# Accelerating High-Performance Classification of Bacterial Proteins Secreted via Non-Classical Pathways: no needing for deepness

**DOI:** 10.1101/2023.01.29.526081

**Authors:** Luiz Gustavo de Sousa Oliveira, Gabriel Chagas Lanes, Anderson Rodrigues dos Santos

## Abstract

Understanding protein secretion pathways is paramount in studying diseases caused by bacteria and their respective treatments. Most such paths must signal ways to identify secretion. However, some proteins, known as non-classical secreted proteins, do not have signaling ways. This study aims to classify such proteins from predictive machine-learning techniques. We collected a set of physical-chemical characteristics of amino acids from the AA index site, bolding known protein motifs, like hydrophobicity. We developed a six-step method (Alignment, Preliminary classification, mean outliers, two Clustering algorithms, and Random choice) to filter data from raw genomes and compose a negative dataset in contrast to a positive dataset of 141 proteins from the literature. Using a conventional Random Forest machine-learning algorithm, we obtained an accuracy of 91% on classifying non-classical secreted proteins in a validation dataset with 14 positive and 92 negative proteins - sensitivity and specificity of 91 and 86%, respectively, performance compared to state of the art for non-classical secretion classification. However, this work’s novelty resides in the fastness of executing non-CSP classification: instead of dozens of seconds to just one second considering a few dozen protein samples or only ten seconds to classify one hundred thousand proteins. Such fastness is more suitable for pan-genomic analyses than current methods without losing accuracy. Therefore, this research has shown that selecting an appropriate descriptors’ set and an expressive training dataset compensates for not using an advanced machine learning algorithm for the secretion by non-classical pathways purpose. Available at https://github.com/santosardr/non-CSPs.

## 1 Introduction

Protein transport can occur from the cytoplasm to other cell compartments, external environments, and other organisms [1]. The process of protein secretion is an important activity for prokaryotic and eukaryotes organisms ([2]. One of the various functions of protein secretion systems is present in disease-causing bacteria. Such pathogenic bacteria use their protein secretion pathways to manipulate the host and establish replication niches [3]. An example is the virulence factors transport through the cytoplasmic membrane toward the host [4] [5]. One of the essential stages in studying diseases and their corresponding treatments is identifying proteins and their secretion pathways [6]. In this sense, learning the routes of protein secretion is paramount in combating various pathologies known to humans. The general secretion pathway (Sec) and the arginine translocation pathway (Tat) are the main routes of protein transport through the cytoplasmic membrane [1]. Proteins secreted by the Sec pathway depend on a signal located in the amino-terminal region of the proteins. This signal comprises twenty amino acids divided into a positively charged N-terminal region, a hydrophobic center, and a polar C-terminal region [7]. As a signal peptide, the proteins secreted by the Tat pathway have a sequence of two arginines in their N-terminal region [8]. In addition, there are pathways present only in gram-negative bacteria, like T1SS, T2SS, T3SS, T4SS, T5SS, and T6SS. Also, only gram-positive bacteria, like SecA1, SecA2, and T7SS, have different signaling regions in secreted proteins [9] [10] [11]. Despite our knowledge about exportation mechanisms, the proportion of proteins predicted as exported is a smaller fraction of a bacterial proteome than the predicted intracellular ones. For instance, the Mycobacteria tuberculosis H37Rv contains about four thousand proteins. A subcellular compartment prediction for proteins within this organism via the SurfG+ software [12] resulted in 13.8% (556) of predicted exoproteins. We have 75.6% or about three thousand proteins classified as cytoplasmic and the remainder (10.46%) as membrane integral. SurfG+ uses the Sec pathway only to decide secreted proteins (5.62%) and several other in-house motifs to differentiate potentially surface-exposed proteins (8.32%), both considered exported proteins. However, some proteins have no signal peptide or apparent identification exportation pattern. Consequently, about 75% of proteins classified by SurfG+ as cytoplasmic are candidates for non-classical secretion. We used to call Non-Classically Secreted Proteins (non-CSPs) molecules primarily sharing only the fact that they locate in the extracellular medium [13]. We know several proteins classified as essential for characterizing infections and diseases and secreted by non-classical pathways [13]. We also use non-classical pathways to produce recombinant proteins in the biotechnological industry [6]. We know prediction methods based on the functional classification of proteins [14] [15] and more recently advanced machine-learning techniques exploring dozens or hundreds of features from protein sequences [16] [17]. Such methods use the analysis of physical-chemical characteristics present in the amino acid sequence on grouping proteins. In this research, we also aimed to classify non-CSPs by selecting physical-chemical propensity indexes as machine-learning descriptors. However, this work’s novelty resides in the fastness of executing non-CSP classification: instead of dozens of seconds to just one second considering a few dozen protein samples or only ten seconds to classify one hundred thousand proteins. Such fastness is more suitable for pan-genomic analyses than current methods without losing accuracy. Besides, we provided a novel methodology to minimize contradictory label assignments to negative training datasets that are broadly applicable.

## 2 Materials AND Methods

We have broken down our methodology into several stages. The first stage consisted of the search for non-CSPs established by the current literature and creating our positive training (141 proteins) and validation (14 proteins) datasets. Next, we search the AA index site for physical-chemical characteristics that classify secretion by non-classical pathways. We chose thirty propensity scales related to hydrophobicity. We used them as descriptors for machine-learning algorithms. We initially created a negative training dataset of 1050 proteins, equally sampled thirty proteins per genome, among thirty-five bacterial genomes downloaded from NCBI, mostly classical secreted ones. We assembled the AA index selected characteristics according to the ARFF format, trained, and tested in all machine-learning classifiers present in the WEKA software [18], capable of dealing with the training data. We compared the results of the different algorithms selecting the best for further analyses. Finally, we recreated the negative training dataset with the full set of proteins from 28 complete genomes to choose a trustable and larger negative training dataset, obtaining a better predictive performance after successfully applying the filters from one to six described below.

### 2.1 Preprocessing

We created an in-house program called valifasta (‘src/valifasta’) to remove punctuation characters, line breaks, and other artifacts occasionally present in downloaded protein sequences. Valifasta aims to minimize probable processing errors from bioinformatics tools that accept only the format of the twenty letters representing the essential amino acids and identification for each protein and ensures each protein will have a unique identification key. We processed all proteins used in this work with valifasta.

### 2.2 Features Search

The affinity of non-CSPs with the plasma membrane contributes to their exportation outside the cellular compartment. The plasma membrane of prokaryotes consists mainly of hydrophilic and hydrophobic regions [19]. We considered the proportion of amino acids with such a characteristic in selecting targets for the training set. In addition, the attributes of basicity and acidity are also essential in the secretion process. Together, these characteristics distinguish four groups of amino acids.

We represented non-polar characteristic amino acids (hydrophobic) by blue color and polar amino acids, i.e., hydrophilic, as purple. We used green to describe essential amino acids and the acid ones in orange. Only these four descriptors cannot correctly classify the main four subcellular sites a protein can be found (cytoplasm, membrane, exposed on the membrane, and outside the cell-secreted by classical pathways). For this reason, we sought new descriptors in the AA index repository [20]. The AA index repository [20] has hundreds of weight propensity indices for each of the twenty essential amino acids. It considers the chemical, physical, and structural characteristics of the amino acids studied in significant sets of organisms, the proportion of three-dimensional alpha-helix structure, the amount of negatively and positively charged amino acids, and types of metals of catalytic sites, among others. We empirically gathered 30 propensity scales from the AA index as our result. The result list in alphabetical order is ACID, AURR980101, AURR980105, AURR980118, BASIC, BEGF750101, BROC820102, CHAM830103, CHAM830104, CHAM830105, FAUJ880111, FAUJ880112, GEIM800103, GEIM800105, LEWP710101, MONM990101, MONM990201, NAKH900102, NAKH900108, NONPOLAR, OOBM850104, PALJ810115, POLAR, PONP800106, QIAN880116, RICJ880107, ROBB760111, ROSM880103, VENT840101, and ZHOH040103. We formatted all our data using the ARFF file format from the Weka software using the above features: thirty original parts from the AA index (file ‘src/propensity.dat’), ninety derived from these thirty original counting amino acids at the beginning, middle, and end of each protein, plus the twenty amino acids. We designed the software ‘src/features.lisp’ in the common language to insert the original, derivative, and twenty amino acid features in all ARFF files, like filtering, training, and validation.

### 2.3 Positive Training Dataset

To create a positive training dataset, we identified and cataloged proteins secreted by non-classical pathways from the recent literature. We acquired 141 positives for non-CSP proteins from the PeNGaRoo article [16].

### 2.4 First Negative Training Dataset

Table 1 depicts the number of genomes by genus used to sample proteins for the training set concerning non-CSPs. We selected these genomes considering pathogenic bacteria to humans. From Table 1, we sampled thirty-five distinct genomes to create our initial negative training dataset. We used the software SurfG+ (folder ‘src/surfg+’) to classify all proteins according to four subcellular locations (cytoplasm, membrane, surface exposed, and secreted). We empirically think of gathering about one thousand proteins to avoid an extreme unbalance compared to the size of the positive training dataset, which contains 141 proteins. Ultimately, we created a negative training dataset containing 1050 proteins, the majority selected as classical secreted ones (file ‘data/surfgplus neg.faa’).

**Table 1.** Genomes by genus creating the negative training set.

| Genus | Number of genomes |
| --- | --- |
| *Acinetobacter* | 3 |
| *Bacteroides* | 1 |
| *Campylobacter* | 2 |
| *Clostridioides* | 1 |
| *Clostridium* | 2 |
| *Corynebacterium* | 1 |
| *Enterobacteriaceae* | 3 |
| *Enterococcus* | 2 |
| *Escherichia* | 6 |
| *Klebsiella* | 1 |
| *Mycobacterium* | 1 |
| *Neisseria* | 1 |
| *Pseudomonas* | 1 |
| *Salmonella* | 2 |
| *Shigella* | 2 |
| *Staphylococcus* | 1 |
| *Stenotrophomonas* | 1 |
| *Streptococcus* | 3 |
| *Vibrio* | 1 |

### 2.5 Second Negative Training Dataset

We created the second negative training dataset from a poll of 28 complete genomes comprising 100238 proteins. We selected these 28 genomes considering pathogenic bacteria to humans, according to Table 1. Compared to the first negative training dataset, we chose a smaller number of genomes keeping at least one species from the genus but more than one for worthy genera like *Acinetobacter, Enterococcus, Campylobacter, Streptococcus*, and *Escherichia*. We designed almost one hundred thousand proteins as our initial candidate training negative dataset. To speed up the filtering process, we created an ARFF file containing all training sequences mapped to the attributes obtained from the AA index, despite most of the entries still being candidates for training. The filtering process was technically reduced to a sed Linux command to remove undesirable lines from the ARFF file.

### 2.6 Validation 1 – my independent dataset (MYIDS)

This is the first set of proteins used to validate the classifiers comprised of 106 proteins. We obtained 14 positives for non-CSP from Wang’s research [21] after removing the 141 included in the positive training dataset. We also considered two proteins with high sequence similarity to the training group. Ninety-two negatives for non-CSP validation came from the UniProt repository [22]. They comprised a set of membrane proteins (integral and partial) exported by classical pathways and cytoplasmic proteins.

### 2.7 Validation 2 – PeNGaRoo original independent dataset (ORIDS)

The PeNGaRoo [16] and ASPIRER [17] software used the same independent validation datasets, ASPIRER, after PeNGaRoo. We directly downloaded 34 negatives and 34 positive validation proteins from the ASPIRER site (files ‘data/pengaroo independent test neg.faa’ and ‘data/pengaroo independent test pos.faa’).

### 2.8 Datasets by Subcellular location

Table 2 lists the subcellular location according to the software SurfG+ (folder ‘src/surfg+’) for all files in the ‘data’ folder. As expected, most of the proteins used by PeNGaRoo were predicted as cytoplasmic. Our work creates the only sets presenting diversity between the four indicated central subcellular locations.

**Table 2.**
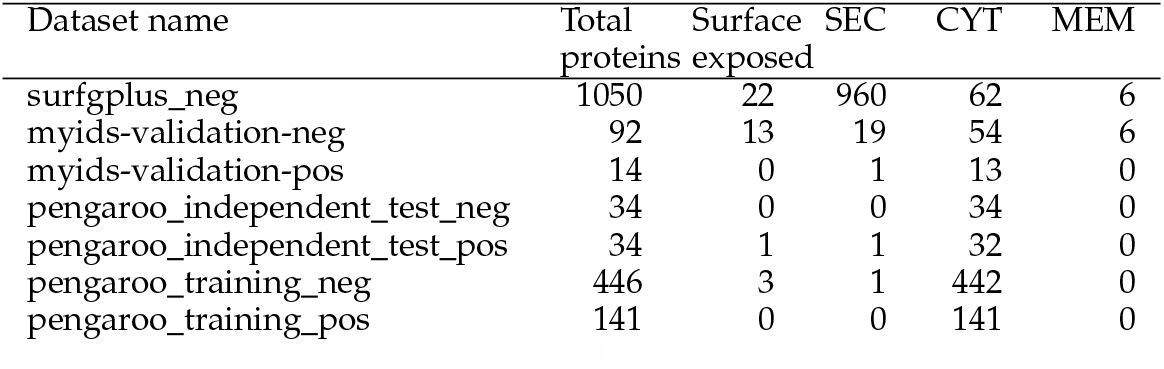
Training and test proteins counting splitting four subcellular locations. Legend: SEC=Secreted, CYT=Cytoplasmic, MEM=Membrane integral.

### 2.9 Data assembly for WEKA

The machine learning software tool we used was WEKA [18]. We calculated the frequency of amino acids of the selected proteins according to the indices chosen feeding the WEKA classifiers algorithms. We used frequencies and the values assigned according to the AA index as weights, thus generating numerical vectors for each protein according to the selected indices. Table 3 exemplifies the weighting of some physical-chemical characteristics used in training algorithms.

**Table 3.** Example of indices (features) and their values for each amino acid.

| Features | A | R | N | D | C | Q | E | G | H | I | L | K | M | F | P | S | T | W | Y | V |
| --- | --- | --- | --- | --- | --- | --- | --- | --- | --- | --- | --- | --- | --- | --- | --- | --- | --- | --- | --- | --- |
| Basic | 0 | 1 | 0 | 0 | 0 | 0 | 0 | 0 | 1 | 0 | 0 | 1 | 0 | 0 | 0 | 0 | 0 | 0 | 0 | 0 |
| Acid | 0 | 0 | 0 | 1 | 0 | 0 | 1 | 0 | 0 | 0 | 0 | 0 | 0 | 0 | 0 | 0 | 0 | 0 | 0 | 0 |
| Polar | 0 | 0 | 1 | 0 | 1 | 1 | 0 | 1 | 0 | 0 | 0 | 0 | 0 | 0 | 0 | 1 | 1 | 0 | 1 | 0 |
| Non-Polar | 1 | 0 | 0 | 0 | 0 | 0 | 0 | 0 | 0 | 1 | 1 | 0 | 1 | 1 | 1 | 0 | 0 | 1 | 0 | 1 |

We also created an in-house program called features, developed in Common Lisp language, to format data in the model required by the WEKA software (‘src/features/features.lisp’). This program processes a MULTIFASTA file, generates a file in CSV format, and counts the values of all amino acids according to selected descriptors. Besides the 30 propensity scales, our software features created three other propensity scales from the original ones. We used our knowledge of the approximate signal peptide size. We counted the amino acids until the border of this maximum size (about 20 AA), creating, for instance, the ACIDINI AA propensity scale for the original ACID propensity scale. We repeat this process at the end of a sequence creating an ACIDEND AA propensity scale. Finally, the AA range between the ACIDINI and ACIDEND constitutes the third derivate AA propensity scale, the ACIDMID. Ultimately, we turned 30 AA propensity scales into 120 AA propensity scales, plus one propensity scale for each amino acid summing 140 attributes or descriptors for each protein. We converted protein sequences into an ARFF file with characteristics signaling secretion or not for non-classical pathways.

### 2.10 Iterative filtering in six steps

We created this iterative filtering process by working with other datasets, like COVID-19. At that time, we perceived a need for filtering false negative entries labeled as negatives, mainly due to the presence of immunoglobulins for COVID-19 in samples of healthy sample donors. Using part of these six steps below, we could produce reliable machine-learning models using the WEKA software capable of high accuracy (unpublished work). Our six steps method to filter false negatives comprises: (i) Remove entries with a significant global Needleman–Wunsch algorithm [23] alignment (*>*90%) against the positive datasets and controls. (ii) Create a prediction model using all the negative and positive training datasets from the PeNGaRoo article [16]. We mapped the 446 negative and 141 positive proteins for non-CSPs to our 140 attributes derived from the AA index. This filter consists of an adapted PeNGaRoo prediction model built by the Weka software (file ‘bin/PeNGaRoo dataset.bin’). (iii) Use the RWeka library in R software for this filtering step, with the file called ‘src/profile.R’. We used the 140 features derived from the AA index in two vectors, one for the positive training dataset and the other for the candidate negative dataset. These vectors contain the mean values of their sets to all features, and we call them positive and negative centroids. After that, we removed entries from the candidate negative dataset possessing a low Pearson correlation (*<*0.95) to the negative centroid and a high Pearson correlation (*>*0.9943) to the positive centroid. We decided on the cut-off values by trial and error according to the results obtained by creating predictive models with the remaining entries. The Linux command sed could not eliminate all lines at once. To accomplish this, we created a script called ‘src/create.sed’ to split the removal into several independent and minor sed delete commands. (iv) Use the RWeka library in R software for this filtering step and the R package “optics dbScan” with the file called ‘src/DBScan.R’. This software tries to create numerically labeled groups of similar elements using the default parameters epsilon=0.9 and minPoints=6. We variate these parameters trying to isolate elements very distinct from the majority and hoping that these are the offending ones for our classifications. (v) Use the RWeka library in R software for this filtering step, with the file called ‘src/Kmeans.R’. A filter like the previous one, using Kmeans, can also pinpoint possible elements to be removed from the training set. (iv) We created a series of scripts in the folder ‘src/semiag/semiagN.bash’ for this filtering step. The N in the script name alludes to the number of elements from the candidate negative training dataset we should draw at once to upgrade our classification results. We run this algorithm recursively and manually, like a genetic algorithm (a semi-genetic algorithm), over the previous better outcome.

## 3 Results

### 3.1 Comparing Classifiers

We compared the results of all classifiers available in the WEKA software. We selected the four best classifiers, highlighting the main developments in Table 4.

**Table 4.** Result of the four best qualifiers for non-CSPs.

| Classifier | Specificity% | Sensitivity% | Accuracy% | False + | False - |
| --- | --- | --- | --- | --- | --- |
| K Star | 73 | 100 | 76 | 25 | 0 |
| Random Forest | 71 | 86 | 73 | 27 | 2 |
| IBk | 66 | 100 | 71 | 31 | 0 |
| Random Committee | 66 | 93 | 70 | 31 | 1 |

Two of the top four classifiers belong to the so-called instance-based learning group. Such classifiers (K Star and IBk) obtained an accuracy of 76 and 73%, respectively. The Instance-based learning method derives from algorithms called K-nearest neighbors (KNN) or classifiers from nearby neighbors. Such algorithms calculate the proximity measure between the analyzed data to generate a classification. Broader used, such algorithms are famous for their ease of implementation, versatility, and high accuracy, which justify the results. To mature our methodology, we chose the algorithm Random Forest [24], the second better algorithm according to our benchmark (Table 4). Besides better results, another argument favoring this choice is the speed of processing the vast datasets containing several thousand entries from this work. In the following subsections, we will describe using Random Forest to refine our methods and decide on better negative training datasets for the task of classifying non-CSP proteins.

### 3.2 Six steps filtering over the MYIDS validation dataset

We applied the filtering methods comprised of the six steps described in Material and Methods over the MYIDS validation dataset. We obtained the following results by filtering action: (i) The alignment filter removed 261 proteins from the initial dataset. However, these are important because they are part of the validation dataset. At the end of this step, 99977 remains in our negative training dataset. (ii) We applied our adapted PeNGaRoo prediction model over the 99977 candidate negative training proteins. As a result, we obtained 90499 classified as non-CSPs capable of proceeding to the next filter. Here we ran a validation getting a completely biased predictor capable of misidentifying all 14 positive validation non-CSPs proteins (sensitivity of 0%). We expected this result because, at this stage, there are probably hundreds of false negatives within the candidate training set. (iii) We obtained an intermediary result over the validation set using 8279 candidates for negative training proteins while searching for cut-off values. This intermediary result gave us specificity, sensibility, and accuracy of 99, 43, and 92%, respectively, for the Random Forest algorithm. However, an unacceptable result considering the low predictive power for non-CSPs. Ultimately, we could remove more than 84 thousand proteins or about 93.2% of our initial candidate negative training dataset. We ended this filtering step with 6152 candidate proteins for the negative training dataset. At this point, validation with the remaining entries culminated in specificity, sensibility, and accuracy of 88, 79, and 87%, respectively, for the Random Forest algorithm (file ‘src/myids-filter2-88-79-87.arff’). This result already overcomes those from Table 4. (iv) Using the default parameters in DBScan (epsilon = 0.9), just a few elements are not included in the single cluster created for our 6152 candidates’ negative training dataset. After we removed the parts outside this single cluster, we could not perceive any improvements in the classification of our validation dataset. After that, we inverted the logic and tried to create the smallest group of clustered elements possible, labeling most entries as non-classified (NA). With parameters epsilon=0.1 and minPoints=6, the algorithm DBScan [25] fragmented the candidate negative dataset to the extreme: 6103 proteins classified as NA’s. Only three sets (14, 21, and 14 proteins) remained. After we removed these 49 proteins, the sensitivity of our classifier was increased from 79 to 86%, followed by a slight decrease in sensitivity from 88 to 86% compared to the previous filtering step (file ‘src/myids-filter3-86-86-86.arff’). We applied the second round of this filter using the parameters epsilon=0.025 and minPoints=3. Now, we got twenty proteins separated into six sets. Removing each batch of proteins in isolation increased the sensitivity at the cost of slightly decreasing specificity. However, instead of eliminating the packs in isolation, we joined them for removal. Two collections removed together produced an actual increase in specificity without diminishing sensitivity. At this point, validation with the 6097 remaining candidate negative entries culminated in specificity, sensibility, and accuracy of 89, 86, and 89%, respectively, for the Random Forest algorithm (file ‘src/myids-filter3-89-86-89.arff’). (v) Working on other data sets, we had the chance to upgrade our results using Kmeans [26]. However, we could not upgrade our results even after splitting the data into 80 clusters for this data. (vi) A random procedure excluding five proteins at once, retraining, and validating allowed us to accomplish specificity, sensitivity, and accuracy of 91, 86, and 91%, respectively (file ‘src/myids-filter5-91-86-91-a.arff’). Another outcome was specificity, sensitivity, and accuracy of 89, 93, and 90%, respectively, after removing three negative entries from the candidate training dataset (file ‘src/myids-filter5-89-93-90-a.arff’). As a result, considering that K Star [27] had a better outcome than Random Forest [24] in Table 4, we trained and validated the performance of this WEKA classifier using the same training data set obtained at step vi. K Star got specificity, sensibility, and accuracy of 67, 100, and 72%, respectively. Despite being consistent with its previous result, it is far worse than our algorithm of choice Random Forest.

### 3.3 Six steps filtering over the ORIDS validation dataset

We also applied the filtering methods comprised of the six steps described in Material and Methods over the ORIDS validation dataset, the original dataset used by PeNGaRoo and ASPIRER. We obtained the following results by filtering action when ORIDS was our driving dataset: (i) The alignment filter removed 86 proteins from the initial dataset. At the end of this step, 100152 remains in our negative training dataset. (ii) We applied our adapted PeNGaRoo prediction model over the 100152 candidate negative training proteins. As a result, we obtained 90718 proteins classified as non-CSPs capable of proceeding to the next filter. (iii) We started from the positive (*>*0.9943) and negative (*<*0.95) cut-off values for the Pearson correlation we used on the MYIDS dataset. Using the starting cut-off values, we got a negative training dataset with 41860 proteins. The validation acquired specificity, sensibility, and accuracy of 91, 50, and 71%, respectively, for the Random Forest algorithm. After experimenting with several other filtering values, we could not obtain a set of proteins that, after removal, could simultaneously keep specificity and sensitivity above 80%. Our last result was a negative training dataset with 5237 proteins, acquiring specificity, sensibility, and accuracy of 44, 88, and 66%, respectively. We opted to pass our initial trial for the next step, hoping we could have a slight decrease in specificity followed by a significant increase in sensitivity. (iv) Using DBScan, we could not avoid the same behavior observed in the previous step: an antagonist behavior between specificity and sensibility. The better result we obtained in this step was a negative training dataset with 36064 proteins, acquiring specificity, sensibility, and accuracy of 79, 62, and 71%, respectively. We obtained this result after running DBScan twice over the initial dataset using epsilon=0.7 and minPoints=6. At this step, we decided to terminate our tries to improve this training dataset because we experienced that further measures could not simultaneously produce significant leaps in specificity and sensitivity. Figure 1 depicts several experiments with the ORIDS training dataset where we can perceive the up and down of specificity and sensitivity but always in different directions.

**Fig. 1.**
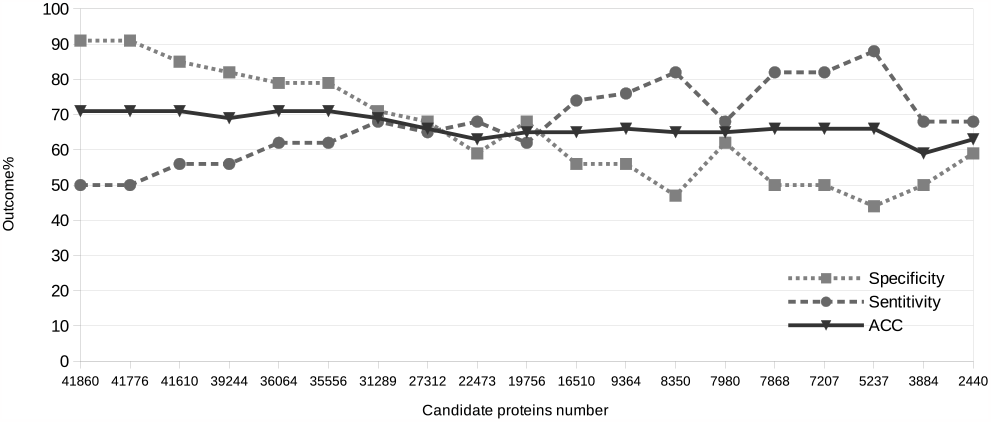
Filtering and testing experiments with ORIDS as the validation dataset.

### 3.4 Comparing models

The training dataset we created using MYIDS (Tests 1 and 3) obtained better performance than those we created using ORIDS (Tests 2 and 4, in Table 5). Besides testing models against our MYIDS (Validation/Set 1), we also tested these against the ORIDS (Validation/Set 2). We gave our models names after Validation 1, the better outcomes. In Validation 2, both our models got performance decreasing (Tests 2 and 4). One can think of the results obtained by our models in tests 2 and 4 discrediting our claimed results. However, our models kept consistently classifying the negative validation dataset from ORIDS with mean specificity and sensitivity of about 86 and 73%, respectively.

**Table 5.** Comparing models produced in this work and state-of-the-art^a^.

| Test | Model | Set | Set<br>size | Spec.<br>% | Sens.<br>% | Acc.<br>% | Time<sup>a</sup><br>(secs) |
| --- | --- | --- | --- | --- | --- | --- | --- |
| 1 | myids5-89-93-90-a | 1 | 106 | 89 | 93 | 90 | 1<sup>b</sup> |
| 2 | myids5-89-93-90-a | 2 | 68 | 85 | 62 | 74 | 1<sup>b</sup> |
| 3 | myids5-91-86-91-a | 1 | 106 | 91 | 86 | 91 | 1<sup>b</sup> |
| 4 | myids5-91-86-91-a | 2 | 68 | 88 | 56 | 72 | 1<sup>b</sup> |
| 5 | PeNGaRoo_dataset | 1 | 106 | 84 | 71 | 82 | 1<sup>b</sup> |
| 6 | PeNGaRoo_dataset | 2 | 68 | 97 | 47 | 72 | 1<sup>b</sup> |
| 7 | PeNGaRoo_website<sup>c</sup> | 1 | 106 | 37 | 100 | 45 | 673 |
| 8 | PeNGaRoo_website<sup>c</sup> | 2 | 68 | 73 | 82 | 77 | 480 |
| 9 | ASPIRER<sup>d</sup> | 1 | 106 | 43 | 92 | 50 | 70+255<sup>e</sup> |
| 10 | ASPIRER<sup>d</sup> | 2 | 68 | 94 | 88 | 91 | 44+80<sup>e</sup> |

Superscript table legends:
<sup>a</sup> We performed the tests and recalculated the metrics for PeNGaRoo and ASPIRER.<sup>b</sup> Considering standalone formatting and classification only.<sup>c</sup> Recommended configuration for quality assurance.<sup>d</sup> -threshold=0.2<sup>e</sup> Creating a Pse-PSSM matrix in a third-party site (POSSUM) + ASPIRER.py processing time.

On the contrary, both state-of-the-art software performed poorly when confronted with our negative validation dataset: PeNGaRoo (test 7) obtained a specificity of 37, and ASPIRER (test 9) reached a specificity of 47%. Low specificity is a measure to be avoided when predicting non-CSPs. Because most of the genome proteins are cytoplasmic, we cannot have the most significant part of proteins submitted to classification labeled as false positives, even with greater specificity. Curiously, even PeNGaRoo classifying its self-created independent negative validation dataset (test 8) is not an exceptional result with a specificity of 73%. We wondered how our chosen AA index descriptors would perform over a training dataset different from ours. Trying to answer such a question, we used the PeNGaRoo training datasets to create a new non-CSPs model. Tests 5 and 6 in Table 5 show these results consistently predicting non-CSPs proteins with 90% of specificity, even with a size 13-fold smaller than our training datasets. For sensitivity, our varied PeNGaRoo model was also steady, despite one modest and other poor results. In Table 5, for tests 5 and 6, the difference from the results presented in Table 4 is a subtle sensitivity instead specificity. To our experience with this work, considering our chosen AA index descriptors, we hypothesized that the unfiltered negative training datasets used to create tests 5 and 6 could shape contradictory signaling to the Random Forest algorithm affecting the capacity of our adapted PeNGaRoo model correctly classify the actual positive proteins. In table 5, there is an unexpected result. Both the predictors of the state-of-the-art and our predictors are biased. The classifiers produced in this work have a bias favoring predicting negatives, while the predictors of the state-of-the-art with a tendency to expect positives. One could argue that the number of elements used to train the models of this work would be responsible for this bias. However, we also obtained above-average results in predicting positives even with this bias for predicting negatives (tests 1 and 3), also an unexpected result considering the tendency due to the overrepresentation of a data class. On average, the specificity and sensitivity of our predictors were 89 and 62%, respectively. On the other hand, state-ofthe-art predictors obtained an average specificity and sensitivity of 69 and 90%, respectively. State-of-the-art predictors, on average, have an almost inverted result compared to the predictors of this study. This bias explains the modest average sensitivity to this work’s classifiers and the ordinary specificity produced by state-of-the-art predictors. A user should assess the models’ performance according to their purposes. Suppose a must high-specificity scenario, the typical scenario on predicting from whole genomes. A researcher cannot get a list of thousands of false positives even if specific non-CPS proteins exist among those. Even our starting predictor in Figure 1 is acceptable in such a picture because we are missing just one for each pair of true positives and pointing out minimal false positives. It happened to Test 6, in Table 5, when we predicted the same independent test dataset used by state-of-the-art software, obtaining 97, 47, and 72% to specificity, sensitivity, and accuracy, respectively. Due to developments like this example, we decided to keep the binary model to our adapted PeNGaRoo dataset within our software repository (file ‘bin/PeNGaRoo dataset.bin’). Another worthy-of-note feature in Table 5 is the time to execute classifications. Our models took about one second to run predictions for both validation sets, considering the time to format (0.6 secs using the script ‘src/createARFF.bash’), classify, and output results (0.3 secs using the script ‘./run.validation1’ and ‘./run.validation2’). Considering a web server, we expect ten seconds as an average expectation for data upload, processing, and mail sent. Other software took hundreds or thousands more times for the same task, even discounting the uploading and mailing times. Another example of our model classification speediness remounts to our crafting efforts to create a negative training dataset. In our second filtering process step, we started with about one hundred thousand candidate proteins (Figure 1). These proteins’ set undergoes the same classification process as Tests 5 and 6 in Table 5. Our adapted PeNGaRoo model took about ten seconds to classify these one hundred thousand proteins, unqualifying about 10% of the proteins as candidates in our ongoing negative training dataset. Besides the speediness, this our adopted model kept consistent with pointing out false positives because 90% of data were not misclassified, as expected, even predicting in a massive dataset.

## 4 Discussion

### 4.1 Why are we using only fourteen proteins as the positive test dataset?

The state-of-the-art works to classify non-CSP proteins present us with less than two hundred proteins known as such [16] [17]. From this total, they used one hundred and forty-one proteins for training purposes, as we also, and thirty-four as a positive validation dataset. We intended to create a novel and more diverse test dataset compared to previous work. Our methodology to create training datasets cannot be used to craft positive ones since we need exportation proof, a feature not effortlessly achieved. Because of that, we took the remaining sixteen proteins as candidates for positive ones in our test dataset. In the end, we filter only fourteen. Note that we also used the validation data set used by the state-of-art against our prediction models. We agree that there are better scenarios than a test set comprising a few cases for a class. However, given that a small number of proteins are broadly known as non-CSP, a workaround has yet to be smoothly achieved.

### 4.2 What could have gone wrong with ORIDS training?

The six steps method for filtering negative training datasets in this work uses the validation dataset to mark out a reasonable candidate dataset. We perform such beaconing by excluding proteins: sequence-similar compared to the validation dataset, classified by a previous training model, correlated to the positive validation dataset, outside the major or inside minor clusters, and finally, the randomized ones. A classification model should perform better without a set of proteins indicated for exclusion. As we can see, the filtering process relays on the validation dataset. Considering our chosen AA propensity index reliance on hydrophobicity and the presence of proteins possessing this trace within the MYIDS validation dataset (Table 2) could explain the reason for a better outcome of the MYIDS over ORIDS. The classification of proteins composing the training and testing datasets in four subcellular locations (Table 2) corroborates the subcellular diversity in the training dataset. In our experience, a single protein entering or leaving the training dataset can increase or decrease the performance of our classifiers. We combined datasets to create other validation sets for training. In our attempts, we kept our negative validation dataset comprised of 92 proteins varying the positive validation data set to 34 (only ORIDS) and 48 (34 from ORIDS and 14 from MYIDS). After the six steps’ filter two and testing against the validation dataset 2, we obtained specificity and sensitivity of 88 and 53%, respectively, using the 92×34 dataset. For the 92×48 dataset, we got specificity and sensitivity of 91 and 44%, respectively. We still got a mean specificity closer than 90% and median sensitivity. Briefing, we could not suddenly benefit from an extended positive validation dataset driving the crafting of our training datasets. We believe that in a scenario where a single protein can change the entire performance of the predictor, the inclusion of dozens of new driving positive validation proteins negatively impacted the performance of our predictors for actual positive instances. We conclude that using our initial positive validation datasets containing fourteen proteins contributed to our best results.

### 4.3 About the better algorithms

In Table 4, the second and fourth-best classifiers (Random Forest and Random Committee) obtained an accuracy of 73 and 70%, respectively. Such classifiers belong to the group of committee algorithms, also called ensemble learning algorithms. This group uses the combination of different models for creating a robust outcome and a committee of experts that meets to deliberate decisions that only one expert would be able to handle. There are numerous ways to craft such model combinations. However, the result shown here specifies the randomization method as the form that generated the highest accuracy values. The Random Forest classifier uses random decision tree sets (decision forests) to generate its analysis. The Random Committee classifier constructs a set of basic classifiers from the same dataset; thus, it uses different seeds of random numbers for each classifier to calculate the average of its predictions. As explained, these methods’ robustness and high performance present their results among the best obtained by such analyses. Considering the algorithms used in this work, the individual differences of each of the best classifiers were responsible for the differences observed between these best results. In Table 4, the delimitation of the four best results in the groups of instance-based and committee-based demonstrates the power of such methodologies in the context of problem-solving from machine learning.

### 4.4 Size and diversity matter

Our first try to reach high specificity and sensitivity in predicting non-CSP proteins used a smaller number of candidate negative training datasets comprised of 1050 proteins. We sampled this initial negative training dataset from several pathogenic bacteria to humans considering four sub-cellular locations (cytoplasm, membrane, surface exposed, and secreted), most (91%) planned as being secreted ones. To our surprise, the set of filters (i to vi) presented in this work was not enough to improve our predictive results when using this negative training dataset (data not shown). At this point, we perceived the need for a more extensive and unbiased negative training dataset concerning subcellular location. We found one solution by extending the initial training dataset from about one thousand to a factor of one hundred without a biased selection of probable subcellular location. Our set of filters could help us create a reasonable machine-learning model since there is enough unbiased data. The state-of-the-art software for non-CSPs designed a negative training dataset composed of 446 proteins. Our first model described in the previous paragraph had about 2-fold in size. Our final negative training solution (MYIDS) used nearly 13-fold in size compared to the state-of-the-art. However, in the early stages, our candidate models used a 94-fold size for the negative training dataset, yet it could perform with 91, 50, and 71% to specificity, sensitivity, and accuracy, respectively (Figure 1). To our expectations, we suspected entirely negatively biased predictions (sensitivity 0%) considering the positive training dataset with only 141 proteins and the negative training dataset 297-fold bigger. Before our three initial filtering processes, we perceived zero sensitivities, but not after that. The Figure 1 data was designed by crafting the ORIDS. We perceived a “To rob from Peter to pay Paul” effect on varying the candidate negative training dataset size when developing the ORIDS and the MYIDS dataset. We found this effect despite the filtering method utilized in the candidate data. The main difference among the filters resides in the fine-tuning of creating lists of candidate deletions. We used this knowledge to focus bulks of removals in the former steps, and graining cuts to the last filters. However, someone can tweak all filters’ parameters to perform massive filtering in all stages.

### 4.5 Future works

As a perspective for continuing this research, we plan to identify these false negatives from the training set using, for instance, literature evidence or sequence and structural alignment methods. Another possible move to excel our predictions could be to figure out other amino acid propensity indexes more prone to the task of non-CSPs classification.

## 5 Conclusion

Our results showed that the specificity and sensitivity are close to 90% with the best models. Using the MYIDS validation dataset demonstrates the power of the chosen physical-chemical characteristics and the six steps filtering process to produce training datasets. We understood that we achieved the objective of obtaining a set of physical-chemical features correctly discriminating non-CSPs and minimizing the error of classifiers filtering the training dataset. We also crafted classification models faster than the current state-of-the-art software and are handy to use in standalone mode without needing third-party software.

## Luiz Gustavo de Sousa Oliveira

Graduated in Computer Science from Centro Universitário do Tri ângulo (2015). Master in Computer Science at the Federal University of Uberl ândia (2020). He was with the COMP2BIO laboratory managed by Dr. Anderson Santos, researching protein secretion by non-classical pathways in bacteria using machine-learning algorithms. He is currently working as a software engineer.

## Gabriel Chagas Lanes

A biologist from the Federal University of Uberlandia (2022). While part of the COMP2BIO laboratory headed by Dr. Anderson Santos, he participated in research activities on COVID diagnosis using machine-learning algorithms. Also, he contributed to the GENPPI software to create ab initio protein networks. His graduation course completion work was about computational modeling of pollination behavior in poricidal anthers in another laboratory. He currently works as a web developer.

## Anderson Rodrigues dos Santos

Ph.D. in Bioinformatics from the Federal University of Minas Gerais (2012). Participated in the assembly and annotation of the first genome project conducted entirely in Minas Gerais (Brazil). Experienced using Computer Science for the assembly and annotation of genomes. Professor at the Federal University of Uberl ândia, Faculty of Computing, since 2013. Part of the post-graduate program guiding master’s students since 2014. Head of the Comp2Bio laboratory, striving to complete student formation in Computer Science with a Biological ground.

